# Human cortical neurons rapidly generated by direct ES cell programming integrate into stroke-injured rat cortex

**DOI:** 10.1101/2024.03.15.585240

**Authors:** Raquel Martinez-Curiel, Mazin Hayj, Oleg Tsupykov, Linda Jansson, Natalia Avaliani, Berta Coll-San Martín, Emanuela Monni, Galyna Skibo, Olle Lindvall, Sara Palma-Tortosa, Zaal Kokaia

**Author notes:** Both authors contributed equally to the manuscript (Sara Palma-Tortosa, Zaal Kokaia).

## Abstract

Stroke is a major cause of long-term disability in adult humans, the neuronal loss leading to motor, sensory, and cognitive impairments. Replacement of dead neurons by intracerebral transplantation of stem cell-derived neurons for reconstruction of injured neuronal networks has potential to become a novel therapeutic strategy to promote functional recovery after stroke. Here we describe a rapid and efficient protocol for the generation of cortical neurons via direct programming of human embryonic stem (hES) cells. Our results show that 7 days overexpression of the transcription factor neurogenin 2 (NGN2) in vitro was enough to generate hES-induced cells with cortical phenotype, as revealed by immunocytochemistry and RT-qPCR, and electrophysiological properties of neurons in an intermediate stage of maturity. At 3 months after translantation into the stroke-injured rat cortex, the hES-induced neurons (hES-iNs) showed immunocytochemical markers of mature layer-specific cortical neurons and sent widespread axonal projections to several areas in both hemispheres of the host brain. Their axons became myelinated and formed synaptic contacts with host neurons, as shown by immunoelectron microscopy. Our findings demonstrate for the first time that direct transcription factor programming of hES cells can efficiently and rapidly produce cortical neurons with capacity to integrate into the stroke-injured brain.

## INTRODUCTION

Ischemic stroke causes loss of neurons and glial cells in a specific area of the brain, which leads to long-term sensory, motor and cognitive impairments in a majority of surviving patients.^1^ Effective treatments to improve function in the chronic phase after the insult are lacking and highly warranted. Stem cell-based approaches hold great promise as potential new therapies for a variety of human brain disorders. The clinical trials with stem cell transplantation performed so far in ischemic stroke have not aimed at cellular replacement. Instead, the modest improvements have been attributed to the so-called bystander effect, i.e., the stem cells’ ability for stimulating plasticity, trophic action and modulating inflammation.^2–5^

Experimental evidence from animal models indicates that stem cell transplantation might induce functional improvement also by replacing the dead cells and reconstructing the stroke-injured brain. We have demonstrated that human long-term neuroepithelial-like stem (lt-NES) cell-derived cortical neurons, produced from induced pluripotent stem (iPS) cells and transplanted into stroke-injured adult rat cortex, establish afferent and efferent morphological and functional connections with host neurons in both hemispheres, improve neurological deficits and regulate motor behavior.^6–8^ The grafted, cortically fated lt-NES cells also give rise to mature oligodendrocytes capable of remyelinating host axons.^9^ In analogy to these findings, human ES cell-derived visual cortical neurons implanted into the neurotoxin-injured mouse visual cortex send axonal projections matching the normal ones and may receive functional synaptic input from the host brain.^10^ Recently, human cerebral organoids placed in large injury cavities in the adult rat visual cortex were found to establish reciprocal connections with the host brain and respond to visual stimuli.^11^ Taken together, these studies provide strong support for the notion that reconstruction of the injured adult brain is possible and raise the possibility that in the future this might be achievable also in a clinical setting. In further support of a possible clinical translation, the cortically fated lt-NES cells also differentiate to mature, functional cortical neurons^12^ and oligodendrocytes ensheathing human axons^9^ when transplanted onto organotypic cultures of adult human cortex. Moreover, the clinical trials with transplantation in Parkinson’s patients show that grafted neurons can survive for many years in the diseased, adult human brain, become functionally integrated and give rise to clinically measurable improvements.^13^

Of major importance for the clinical translation of these findings will be to identify the best stem cell source for brain repair, e.g., in patients with ischemic stroke. Stem cells aiming for cell replacement in a clinical transplantation setting must be of human origin. The potentially best stem cell source will be determined by its various properties as assessed in preclinical studies: lack of tumorigenicity, morphological and functional characteristics after grafting in animal models, ability to generate the specific subtypes of neurons which need to be replaced and to generate other cells such as myelinating oligodendrocytes. Moreover, the preclinical protocol should be easily translatable into a clinically applicable one.

Here we have tested a protocol to differentiate human ES cells via direct transcription factor programming to layer-specific functional cortical neurons (hES-iNs).^14^ The objectives were three-fold: First, to characterize *in vitro* the hES-iNs prior to transplantation in a rat model of cortical ischemic stroke; Second, to analyze the survival, phenotype, and projections of the grafted neurons; Third, to determine whether the graft-derived axons form synaptic contacts and become myelinated by oligodendrocytes in the host.

## MATERIALS AND METHODS

### hES-iN generation

hES-iNs were generated as previously described.^14^ Briefly, WAO1 (H1) hES cells were cultured in feeder-free conditions on Matrigel-coated 6 well plates in mTeSR medium. After reaching 80% confluence, cells were dissociated with Accutase and 5×10^5^ cells per well re-plated on Matrigel-coated 6 well plates in mTeSR medium supplemented with Rock Inhibitor (10 μM, Y27632, Selleckchem). 24h after seeding, the hES cells were transduced with 1 μL of TetO-Ngn2-T2A-Puro lentivirus and 1 μL of FUW-M2rtTA lentivirus. One day after infection (*day 0*), the mTeSR medium was replaced with an induction medium (DMEM/F12 supplemented with N2 and B27), where doxycycline (2.5 μg /mL, Sigma) was added to induce NGN2 expression. Doxycycline was kept in the medium until the end of the experiment. On *day 1*, a 3-day puromycin (1.25 μg/mL) selection period was started. On *day 6*, cells were detached with Accutase and collected for RT-qPCR. On day 7 cells were resuspended in a) cytocon buffer supplemented with doxycycline for transplantation or b) induction medium to reseed in matrigel-coated coverslips for immunostainings or electrophysiological recordings.

### Animals

Six athymic nude rats (Crl: NIH-Foxn1 RNU), 8 weeks of age, were purchased from Charles River Laboratories. The animals were housed in individually ventilated cages under standard temperature and humidity conditions and a 12-hour light/dark cycle with free access to food and water. All procedures were conducted in accordance with European Union Directive 2010/ 63/EU and approved by the ethical committee for the use of laboratory animals at Lund University and the Swedish Department of Agriculture (Dnr. 5.8.18-07222/2021 M68-16).

### Distal middle cerebral artery occlusion and cell transplantation

Focal ischemic injury was induced in the somatosensory cortex by distal middle cerebral artery occlusion (dMCAO) as described previously with some modifications.^15,16^ Briefly, animals were anesthetized with isoflurane (3.0% induction; 1.5% maintenance) mixed with air, exposing the temporal bone. A craniotomy of 3 mm was made, the *dura mater* was carefully opened, and the cortical branch of the middle cerebral artery was ligated permanently by suture, cauterized, and cut. Both common carotid arteries were isolated and ligated for 30 min. After releasing common carotid arteries, surgical wounds were closed.

Intracortical transplantation of hES-iNs was performed stereotactically 48 h after dMCAO as described previously.^8^ Briefly, on the day of surgery, programmed hES-iNs on day 7 were resuspended to a final concentration of 100.000 cells/μL in cytocon buffer. A volume of 1 μl was injected in three sites at the following coordinates (from bregma and brain surface): anterior/posterior: +1.5 mm; medial/lateral: +2.0 mm; dorsal/ventral: −2.0 mm; and anterior/posterior: +0.5 mm; medial/lateral: +1.5 mm; dorsal/ventral: −2.0 mm and anterior/posterior: +0.5 mm; medial/lateral: +2.5 mm; dorsal/ventral: −2.5 mm.

### Immunocytochemistry and immunohistochemistry

hES-iNs plated on glass coverslips were fixed on day 8 in 4% formaldehyde (Sigma) for 20 min at room temperature. Cells were permeabilized with 0.025% Triton X-100 in 0.1 M potassium phosphate buffered saline (KPBS) and blocked with 5% of normal donkey serum (NDS) for 45 min. Primary antibodies (**Table S1**) were diluted in blocking solution and applied overnight at +4°C followed by 3 rinses with KPBS. Fluorophore-conjugated secondary antibodies (**Table S1**) (1:500, Jackson Laboratories) diluted in blocking solution were applied for 2 h at room temperature. Afterward, cells were rinsed 3 times with KPBS, and nuclei were stained with Hoechst (Molecular Probes) for 10 min in KPBS at room temperature. Stained glass coverslips were mounted on slides with Dabco (Sigma) mounting media.

Immunohistochemistry in rat slices was done as follows: the stored sections were rinsed 3 times with KPBS and incubated in a blocking solution for 1 h (0.25 Triton X-100 in 0.1 M KPBS [TKPBS] with 10% NDS). The rest of the procedure for cell cultures follows the procedure described above.

Some stainings required antigen retrieval (see **Table S1**) before the permeabilization step. Cells and rat sections were incubated with sodium citrate pH 6.0 Tween 0.05% for 30 min at +65°C.

### Microscopical analysis and quantification

*In vitro* quantifications were performed using 20x images taken using a confocal microscope (LSM 780, Zeiss, Germany). Using Image J software, several positive cells were counted by sampling various regions from different coverslips. The total number of cells in each region of interest was counted using Hoechst staining.

Overview images of rat brain slices stained with STEM101, NeuN, DCX, and Ki67 were taken using a Virtual Slide Scanning System (VS-120-S6-W, Olympus, Germany).

The volume of infarction was measured in sections stained with NeuN. The intact area was determined by cells in both the ipsilateral and contralateral hemispheres, marked out, and measured using Image J software. The infarcted area was determined by subtracting the non-lesioned (NeuN-stained) area in the damaged hemisphere from the corresponding location in the contralateral hemisphere. The lesion volume was calculated by multiplying the infarcted area by the thickness and spacing between sections (300 μm).

The graft area was determined using sections stained with STEM101. The immature and mature regions within the graft were determined by outlining areas positive for DCX and NeuN, respectively. Co-expression was assessed via confocal microscopy by observing the overlap of the two chosen markers within the same plane and area.

Evaluation of areas reached by hES-iNs-derived fibers was performed using a Virtual Slide Scanning System (VS-120-S6-W, Olympus, Germany). Fiber density was assessed semi-quantitatively in 10 μm thick maximum intensity projection confocal images captured with a 63X objective. Three to five images were analyzed for each area.

### RT-qPCR

RNA extraction from hES-iN at D6 of programming was performed with the RNeasy Mini Kit (QIAGEN) following the protocol described by the manufacturer. RNA purity and concentration were determined using a NanoDrop spectrophotometer (ND-1000). RNA (1 mg) was used for cDNA synthesis with qScript cDNA SuperMix (QuantaBio). TaqMan probes (Thermo Fisher Scientific; NANOG Hs02387400_g1, HS00300164_s1; DCX, Hs00167057_m1; BIIITUB, Hs00964963_g1; MAP2, Hs00258900_g1; POU3F2, Hs00271595_s1; and TBR1 Hs00232429_m1) were used and RT-qPCR was run in triplicate samples on a QuantStudio™ 1 Real-Time PCR System (ThermoFisher Scientific) with GAPDH as the housekeeping gene (HS02786624_g1)

### Electrophysiology

Electrophysiological properties of hES-iNs were recorded on day 7. For whole-cell patch-clamp recordings, coverslips were transferred to the recording chamber and constantly perfused with carbonated artificial cerebrospinal fluid (ACSF) containing (in mM): 119 NaCl, 2.5 KCl, 1.3 MgSO4, 2.5 CaCl2, 26 NaHCO3, 1.25 NaH2PO4, and 11 glucose (pH ∼7.4, osmolarity ∼305 mOsm). Patch pipette was filled with an internal solution containing (in mM): Kgluconate 122.5, KCl 17.5, NaCl 8, KOH-HEPES 10, KOH-EGTA 0.2, MgATP 2, and Na3GTP 0.3 (pH ∼7.2, 295 mOsm). The average pipette tip resistance was ∼4-5 MW, and recordings were done at 32° C. Pipette current was corrected online before gigaseal formation. In contrast, fast capacitive currents were compensated for during cell-attached configuration. All recordings were done using a HEKA double patch EPC10 amplifier (HEKA Elektronik, Lambrecht, Germany) and sampled at 10 KHz. PatchMaster software was used for data acquisition, and FitMaster and IgorPro for offline analysis.

Resting membrane potential (RMP) was measured in current clamp mode at 0 pA immediately after establishing whole-cell configuration. Input resistance (Ri) was calculated from a 5 mV pulse and monitored throughout the experiment. The ability to generate an action potential (AP) was determined by 500 ms square depolarizing current step injections at RMP, with 10 pA increments and ramp injection of 1 s depolarizing current, which was also used to determine the action potential threshold. AP amplitude was measured from threshold to peak, the half AP amplitude width was defined as the time between the rising and decaying phase of the AP measured at half the amplitude of the AP, and the afterhyperpolarization (AHP) amplitude was determined as the difference between the AHP peak and the AP threshold. Whole-cell sodium and potassium currents were observed in voltage-clamp mode at a holding potential of −70 mV, and 200 ms voltage steps were delivered in 10 mV increments. To minimize recording errors, series resistance was compensated to 60-80%, and the recording quality was monitored with a test pulse throughout the recording.

### iDISCO

iDISCO tissue clearing was performed according to Renier and collaborators ^17^ with some modifications. Briefly, animals were perfused and post-fixed in 2% PFA for one hour, then kept in PBS at 4°C shortly before the protocol started. Brains were washed three times in PBS for 30 min at RT. Afterward, alcohol dehydration was performed in methanol/PBS with progressively higher methanol concentrations. The concentrations of methanol used were 20, 40, 60, 80, and 100%, and each dehydration step was performed for 30 min at RT. A last 100% methanol incubation was done for 1 h. The samples were then incubated overnight in 66% dichloromethane (DCM) and 33% methanol at RT, washed twice in 100% methanol for 1 h, and then kept overnight at 4°C. On day 3, brains were incubated in freshly prepared 5% hydrogen peroxide overnight at 4 °C. Following overnight incubation, the samples were rehydrated in methanol/0,2%TritonX-100 in PBS with progressively lower methanol concentration. The concentrations of methanol used were 80, 60, 40, and 20% methanol, and finally, 0,2% TritonX-100+PBS, with each rehydration step being performed for 30 min. After that, two washes for one hour were done in 0,05% sodium azide and 0,2% TritonX-100 in PBS at RT with shaking. Following the washings, the samples were incubated in 0,05% sodium azide, 20% dimethyl sulfoxide (DMSO), 0,2% TritonX-100, and 0.3 M glycine in PBS for 5 days at 37 °C. On day 9, two washings with 0,05% sodium azide, 0,2% TritonX-100, and 10ug/mL heparin in PBS were done for 30 min at RT. Samples were then incubated with 0,05% sodium azide, 0,2% TritonX-100, 10% DMSO, and 6% of normal donkey serum (NDS) in PBS for 10 days at 37°C. On day 19, samples were washed two times for half an hour with 0,05% sodium azide, 0,2% TritonX-100, and 10 mg/mL heparin in PBS at RT. After the washings, brains were incubated in a filtered solution containing the primary antibodies SC 101 (Mouse 1:500, Stem Cells Inc) and SC 121 (Mouse 1:500, Stem Cells Inc) diluted in 0.05% sodium azide, 5% DMSO, and 3& NDS in 0.2% Tween- 20 and 10 mg /mL heparin in PBS, for 10 days at 37°C. On day 29, the samples were washed 16 times for 25 min with 0.05% sodium azide, 0.2% Tween 20, and 10 mg /mL at RT. Following the washings, the brains were incubated in a filtered solution containing the secondary antibodies (Cy5, donkey anti-mouse, 1:500) diluted in 0.05% sodium azide and 3% NDS in 0.2 Tween-20 and 10 mg/mL heparin in PBS for 10 days at 37 °C. On day 39, the samples were washed 16 times for 15 min with 0.05% sodium azide, 0.2% Tween 20, and 10 mg /mL at RT. After washing, a methanol/PBS dehydration process was performed, and the samples were incubated overnight with DCM and methanol. On day 40, brains were incubated twice in DCM for 25 min and then in dibenzylether (DBE) until the samples became clear. Before microscope analysis, samples were stored in DBE at RT with no light or air exposition.

Tissue cleared samples were imaged in a sagittal orientation using a sCMOS-5.5-CL3 camera equipped light sheet microscope (Ultramicroscope II, LaVision Biotec) with a 2×/0.5 objective lens (MVPLAPO 2×) with a 6-mm working distance dipping cap. All imaging used the Imspector-Pro219 software and scanned continuously with a step size of 10 μm at 3.2× magnification (7988 × 9472 pixels). Post imaging visualization utilized the Arivis Vision 4D v.2.12.3 software.

### Immuno-electron microscopy (iEM)

Rats were deeply anesthetized with pentobarbital and transcardially perfused with 0.1 M phosphate buffer saline (PBS) followed by ice-cold 2% paraformaldehyde, containing 0.2% glutaraldehyde, in 0.1 M PBS, pH 7.4. Brains were removed and then washed in 0.1 M PBS. Frontal 100 μm sections of the whole brain were cut on a Vibratome VT1000A (Leica, Germany). Sections were postfixed in 1% osmium tetroxide in 0.1M PBS, dehydrated in a graded series of ethanol and propylene oxide, and flat-embedded in Epon. Ultrathin sections were cut with a diamond knife. For post-embedding immunogold labeling of STEM121, ultrathin sections were incubated overnight in primary mouse anti-STEM121 antibody (1:500, Stem Cells) at +4°C. A secondary antibody (goat anti-mouse IgG conjugated to 15 nm colloidal gold; Electron Microscopy Sciences) diluted 1:20 in 0.1% BSA in PBS was added for 1.5 h, then washed with PBS. Sections were then fixed with 2% glutaraldehyde and washed with PBS, followed by dH_2_O. Sections were stained with uranyl acetate and lead citrate. Ultrathin sections were examined and photographed using a transmission electron microscope JEM-100CX (JEOL, Japan).

### Statistical analysis

Statistical analysis was performed using Prism 9 software (GraphPad). An unpaired t test was used when data were normally distributed, whereas a Mann-Whitney U test was used when data did not pass the normality test. Significance was set at p < 0.05. Data are mean ± SEM.

## RESULTS

### Directly programmed human ES cells express markers for mature cortical neurons already after 7 days of induction *in vitro*

In our previous study^14^, we found that the induction of NGN2 gene expression in hES cells gives rise to neuron-like cells by the end of the first week *in vitro.* After two weeks, these cells expressed the mature neuronal marker MAP2 and showed the morphology of mature neurons, exhibiting pyramidal-shape with rich arborization comparable to human adult cortical neurons. Here we assessed in more detail the level of differentiation of the generated hES-iNs after 7 days *in vitro* using the same protocol. Protein and gene expression of different neuronal markers was analyzed as well as the electrophysiological properties of the cells.

The hES-iNs were immunostained for both immature and mature neuronal markers. We observed that 64.4±3.4% of the programmed cells expressed the neuroblast and immature neuronal marker doublecortin (DCX) after 7 days *in vitro* while 54.9±4.3% of the cells expressed the immature neuronal marker Tuj1. In this context, all Tuj1 immunopositive cells expressed DCX (**Figure 1A**). To visualize the generation of mature neurons we stained for NeuN. This marker was expressed in 56.7±4.8 % of hES-iNs and also showed colocalization with DCX (**Figure 1A and B**). Regarding the proliferation state of the cell culture at 7 days of programming, we observed that Ki67 was expressed only in 6.1±1.7% of the cells (**Figure S1A**). Also, we found no expression of the pluripotency marker Nanog and just few cells (<1%) were immunopositive for the multipotency marker Sox2 (data not shown), suggesting the presence of a small population of neural stem cells. Taken together, the data based on the expression of neuronal markers provide evidence that about a 60% of the hES-iNs are at an intermediate stage between immature and mature neurons following 7 days of differentiation.

**Figure 1.**
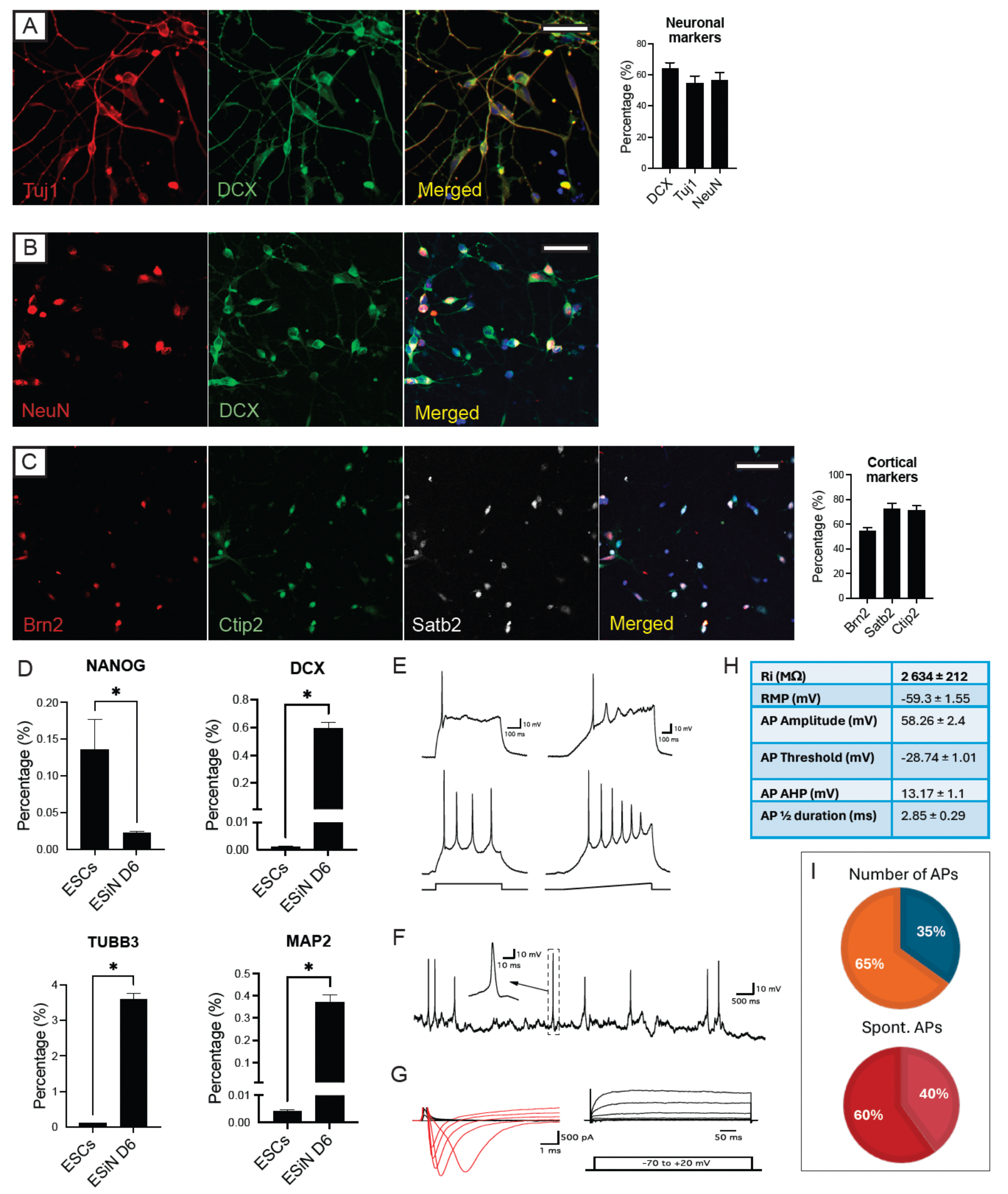
Overexpression of NGN2 in hES cells generate immature and mature cortical neurons after 7 days of programming *in vitro.* **A-C,** Confocal images and quantifications of hES-iNs at day 7 of programming showing expression and percentage of the neuronal marker Tuj1 (**A**), the neuronal progenitor marker DCX (**A, B**), the mature neuronal marker NeuN (**A,B**); and upper and deep cortical layer markers Brn2 and Satb2; and Ctip2, respectively (**C**). Scale bar, 50 µm. **D**, RT-qPCR analysis of gene expression (*NANOG*, *DCX*, *TUBB3* and *MAP2*) in ES cells and hES-iNs after 6 days of programming. Data are presented as mean ± SEM. Significance was set as p < 0.05. **E**, Whole-cell patch-clamp recordings from hES-iNs at day 7. Voltage traces illustrate ability of the cells to generate single (upper traces) or multiple (lower traces) APs during current step (left, 30 and 20 pA, respectively) or ramp (right, 0-50 pA) injection. **F**, Voltage trace illustrating ability of a portion of cells to spontaneously generate APs at RMP. **G**, Current traces illustrating the inward sodium (red) and outward sustained potassium (black) currents observed during voltage steps depolarisation from −70 mV with 10 mV steps. **H**, Table summarising passive membrane properties (Input resistance - Ri and RMP) and AP characteristics of the recorded cells (n=23). **I**, Pie-charts showing a ratio of the cells with (60%), or without (40%) spontaneous APs (n=15) and of the cells with single (35%), or multiple (65%) APs upon depolarization challenge (n=23).

Regarding the neuronal phenotype, 91.9±2.1% of the hES-iNs were excitatory, expressing the glutamatergic marker KGA (**Figure S1B**). In contrast, no cells expressed GAD65/67+ and were of inhibitory phenotype (data not shown). The hES-iNs expressed cortical markers characteristic of both upper layers, such as Brn2 (54.6±2.7%) and Satb2 (72.2±4.5%), and deep layers, e.g., Ctip2 (71.1±3.8%) (**Figure 1C**). In 52 ± 3.4% of the programmed cells co-expression of both, upper and deep cortical layer markers was found. Regarding the generation of other neural cell types, no expression of the astrocyte or oligodendrocyte markers GFAP and Olig2, respectively, was found (data not shown). The results obtained using immunocytochemistry were in line with the gene expression analyzed by RT-qPCR at day 6 of programming. We found a decrease in the expression of the pluripotency marker NANOG and an increase of the neuronal markers DCX, TUBB3 (Tuj1) and MAP2 in cells programmed for 6 days compared to ES cells (**Figure 1D).** We also observed in increased expression of specific cortical markers such as POU3F2 (Brn2; a marker of upper layers) and TBR1 (a marker of deep layers) compared to non-programmed ES cells (**Figure S1C**).

We then performed whole-cell patch-clamp recordings to assess the electrophysiological properties of the hES-iNs at 7 days. The recorded cells already had all the basic properties of functional neurons, i.e., the ability to fire Action Potentials (APs) (**Figure 1E-F**) and expressing fast inward sodium and outward sustained potassium currents (**Figure 1G**). The average resting membrane potential (RMP) of the recorded cells was −59.3 ± 1.55 mV with a majority of the cells (65%, 15 out of 23 in total) being able to generate multiple APs upon step, or ramp current injection, while a smaller portion (35%, 8 out of 23 in total) only fired a single AP (**Figure 1E and I**). These observations indicate the occurrence of more and less mature neuronal populations simultaneously. We also observed spontaneous AP firing at RMP in a good portion (40%, 6 out of 15 in total) of the neurons with multiple APs (**Figure 1F and I**). When characterizing the basic parameters of APs, we observed that on average, there were fast, high amplitude APs with kinetics corresponding to a well-functioning neuron (Fig. 1 H; AP amplitude = 58.26 ± 2.4 mV; AP threshold = −28.74 ± 1.01; AP half duration = 2.85 ± 0.29 ms and afterhyperpolarization potential (AHP) = 13.17 ± 1.1 mV). However, when looking at passive membrane properties, the average input resistance (Ri) was 2 634 ± 212 MΩ, which is rather high but not unusual for newly programmed and still maturing cells with small soma and not very complex arborization. No spontaneous synaptic activity was detected in either of the recorded 23 cells (from 7 coverslips of 2 different programming) at this timepoint. To summarize, the electrophysiological analysis indicates that the hES-iNs, although they have acquired certain neuronal characteristics, are still young and not synaptically integrated at 7 days.

### Directly programmed human ES cells differentiate into layer-specific cortical neurons after transplantation into stroke-injured rat cortex

We have previously shown that NGN2-induced hES-iNs placed onto slices of adult human cortex had survived and differentiated to functional neurons with extended neurites at 4 weeks after transplantation.^14^ Also, some hES-iNs expressed the upper layer cortical marker Satb2. Here we wanted to determine the survival capacity and behaviour of the NGN2-induced hES-iNs after intracortical transplantation in a rat model of cortical ischemic stroke.

Rats were subjected to dMCAO followed by MRI 24h later to confirm the location of the lesion. 48h later, hES-iNs programmed for 7 days were transplanted in close proximity to the damaged cerebral cortex. Animals were sacrificed 3 months thereafter. As observed 24 hours after stroke, the damaged area was restricted to the somatosensory cortex in all animals after 3 months, with variability in infarct sizes and an average of 8.6±3.1 mm^3^.

The vast majority of the grafted cells, identified with the human nuclear marker STEM101, were located close to the lesion covering both somatosensory cortex and part of the M1 motor cortex. Only in the larger grafts a substantial number of human cells were also found in the corpus callosum (**Figure 2SA**). We observed that grafted cells expressed marker of both, immature and mature neurons (DCX and NeuN positive cells, respectively). The distribution of cells expressing these two markers within the transplant core was heterogeneous. In general terms, some areas only expressed DCX (39.98 ± 4.3% of the graft area), while others solely exhibited NeuN expression (35.29 ± 3.1%) (**Figure 2A**). In other areas both markers were found in close proximity within the graft (**Figure 2B**). A minority of grafted cells co-expressed DCX and NeuN, suggesting that they were, at this time point after the transplantation, in a transitional stage towards maturity (**Figure S2B**). Quantification of double positive cells was not possible due to the large number of cells in the graft and the cytoplasmic nature of DCX staining.

**Figure 2.**
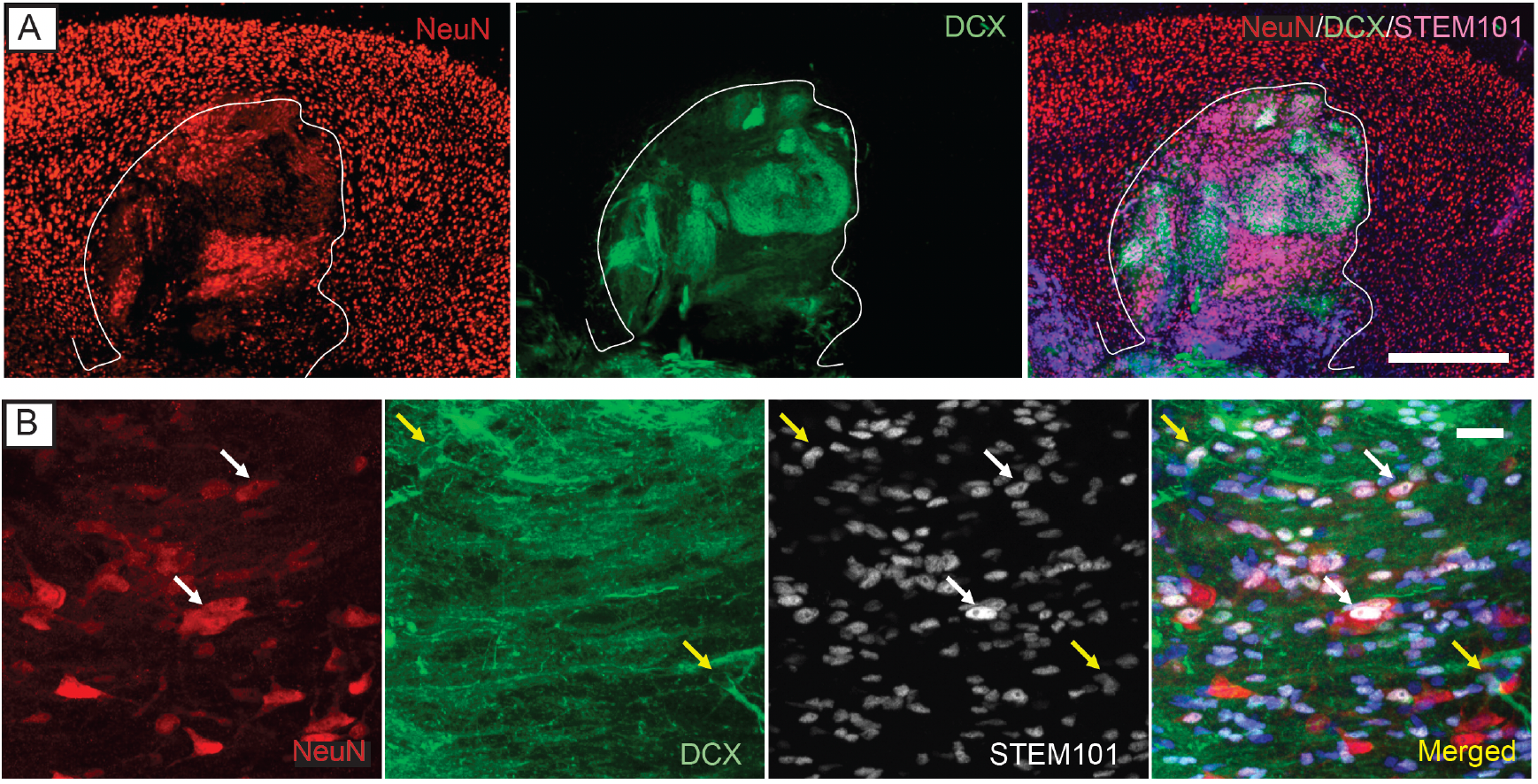
Grafted hES-iNs express markers of mature and immature neurons 3 months after transplantation into rat injured somatosensory cortex. **A-B**, Confocal images showing neuronal progenitors (DCX, green) and mature neurons (NeuN, red) in the transplant core (human nuclear marker, STEM101+ cells). **A**, Overview of grafted hES-iNs expressing NeuN and DCX. Scale bar, 500 µm. **B**, High magnification confocal images of colocalization of NeuN (white arrows) and DCX (yellow arrows) with the human nuclear marker STEM101. Nuclear staining (Ho: Hoechst, blue) is included in merged panel. Scale bar, 20 µm.

To assess the proliferative status of the graft 3 months after the transplantation, we stained for the proliferation marker Ki67 and colocalized with the human marker STEM101. We found sparsely expression of Ki67 within the vast majority the graft (**Figure S2C**) and a higher expression in less mature areas characterized by rosette morphology (**Figure S2D**). Regarding the generation of other neural cell types derived from the transplantation, we found that less than 1% of the cells were graft-derived astrocytes (stained with STEM123 marker, human specific GFAP) and around 1% was graft-derived oligodendrocytes (Olig2-STEM101 positive cells) (**Figure S3A and B**).

We next analysed if the grafted cells expressed layer-specific cortical markers that recapitulate the architecture of the cerebral cortex. Within the transplant core, individual hES-iNs (STEM101^+^ cells) were immunopositive for upper cortical layer markers such as Brn2 (layers II-III, **Figure 3A**, overview and higher magnification images) and Satb2 (II to V, **Figure 3B**). In addition, we found that grafted cells expressed deep cortical layer markers, like Ctip2 (V and VI, **Figure 3C**) and Tbr1 (layer V and VI, **Figure 3D**)). Interestingly, cells expressing the different cortical markers were disposed in a layering pattern as shown in **Figure 3A-D**, spatially separated within the graft. In contrast to the *in vitro* observations, we did not find co-expression of markers for upper and deep cortical layers in individual grafted cells. In this context, we also found immature neurons co-expressing DCX and the cortical marker Satb2 within the core of the graft (**Figure S2E**).

**Figure 3.**
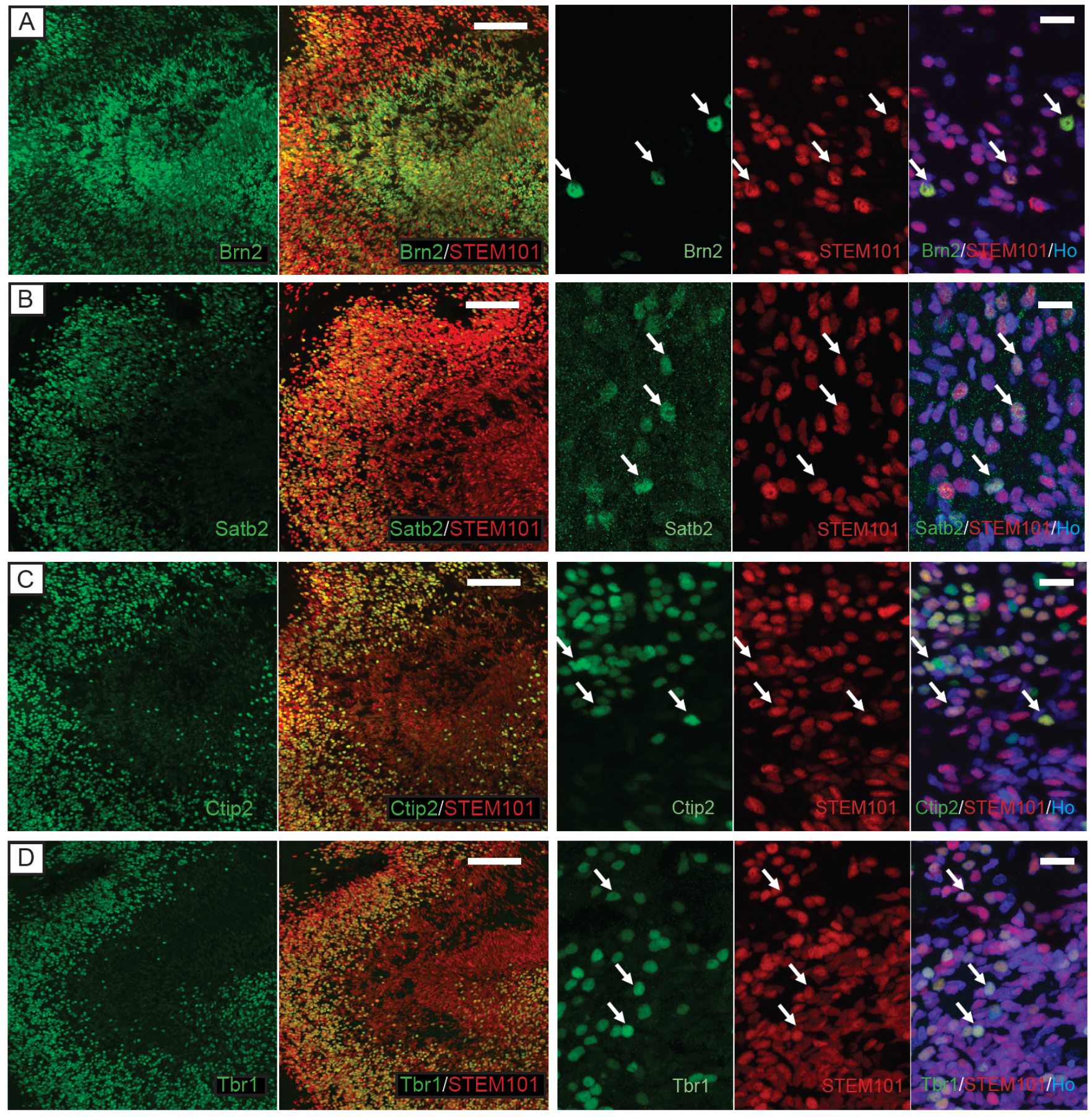
Grafted hES-iN differentiate into layer-specific cortical neurons 3 months after transplantation into rat injured somatosensory cortex. **A-D**, Confocal images showing co-expression of human nuclear marker (STEM101+ cells, red) and the upper cortical layer markers Brn2 (**A**) and Satb2 (**B**); and deep cortical layer markers Ctip2 (**C**) and Tbr1 (**D**) in the core of transplantation. Arrows indicate colocalization. Scale bar low magnification, 500 µm. Scale bar high magnification, 20 µm.

### Grafted, directly programmed hES cells project widely, become myelinated and form synapses with host neurons in stroke-injured rat brain

After characterizing the transplant core, we analyzed the projections of the grafted hES-iNs to the different regions of the brain 3 months after cortical stroke (**Figure 4** and **Video S4**) A large number of graft-derived fibers (stained by STEM121, a human cytoplasmic marker) reached the peri-infarct area (**Figure 4A**) and a substantial number was found in the ipsilateral frontal and somatosensory cortices. Also, a high density of fibers was distributed throughout the corpus callosum (**Figure 4B**) and few of them reached the contralateral somatosensory cortex (**Figure 4C**). Moreover, few fibers were located in the caudate-putamen, septum and internal capsule.

**Figure 4.**
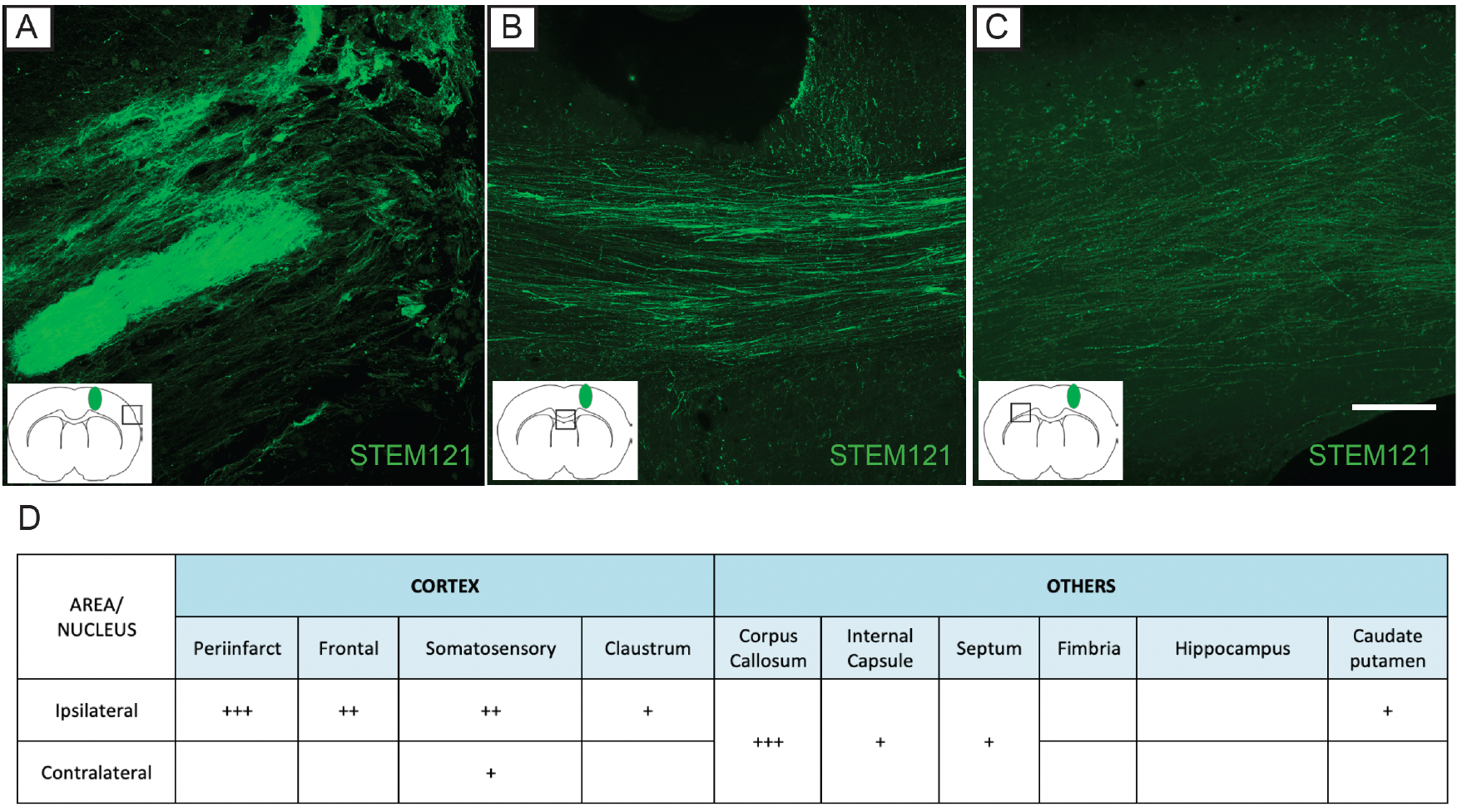
Intracortical grafts of hES-iN send widespread axonal projections in stroke-injured rat brain. **A-C**, Presence of fibers expressing the human cytoplasmic marker STEM121 in different areas of the rat brain: peri-infarct area (**A**), corpus callosum (**B**) and contralateral somatosensory cortex (**C**). Scale bar, 100 µm. **D**. Semiquantitative representation of the density of STEM121+ fibers derived from the graft: +, low density 1 to 5 fibers; ++ medium density up to 50 fibers; and +++, high density more than 50 fibers.

To determine if the axons derived from the grafted hES-iN cells had become myelinated and established synaptic connections with host cells, we performed post-embedding immunogold labeling of STEM121 on ultrathin sections. Immunoelectron microscopy (iEM) analysis demonstrated the presence of individual 15 nm gold particles in STEM121-positive axons and axon terminals. We observed STEM121^+^ hES-iN-derived axons in the corpus callosum and in the ipsilateral and contralateral cortices (**Figure 5A-C**). The human STEM121^+^ axons exhibited ultrastructural features similar to those of host axons, such as microtubules, neurofilaments, and mitochondria, and were enclosed by compact myelin sheaths originating from STEM121-host oligodendrocytes (**Figure 5A-C**).

**Figure 5.**
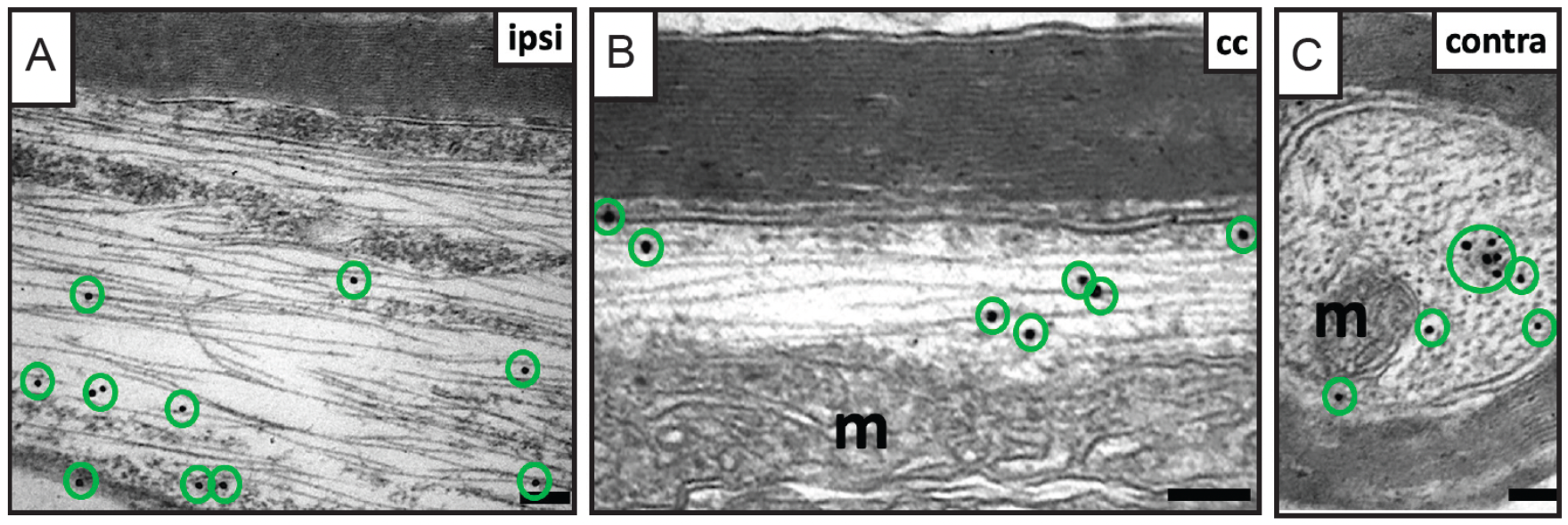
Grafted hES-iN derived-axons are myelinated by host oligodendrocytes in the stroke-injured rat brain. Representative iEM images of STEM121^+^-hES-iN derived-axons myelinated by host oligodendrocytes in corpus callosum (cc, **B**), ipsilateral (ipsi, **A**) and contralateral (contra, **C**) somatosensory cortex. Note: Gold particles are marked with green circles. Scale bars, 0.1 μm.

The iEM analysis also showed that the hES-iN-derived axon terminals had established characteristic synapses with host dendritic spines in the ipsi-(**Figure 6A**) and contralateral (**Figure 6B**) cortex. These STEM121^+^ synaptic contacts had vesicles in the presynaptic terminal, a synaptic cleft, and a postsynaptic membrane with postsynaptic densities. The vast majority (91.7%) of postsynaptic densities were continuous nonperforated (**Figure 6B**), and only 8.3% were perforated (**Figure 6A**). Most (89.4%) STEM121^+^ synaptic contacts were axodendritic and displayed the ultrastructural characteristics of asymmetric excitatory/glutamatergic synapses, e.g., prominent postsynaptic density, a wide synaptic cleft, and spherical synaptic vesicles. The STEM121+ presynaptic terminals had docked synaptic vesicles at the presynaptic membrane (**Figure 6**), indicating the functional activity of synapses.^18^

**Figure 6.**
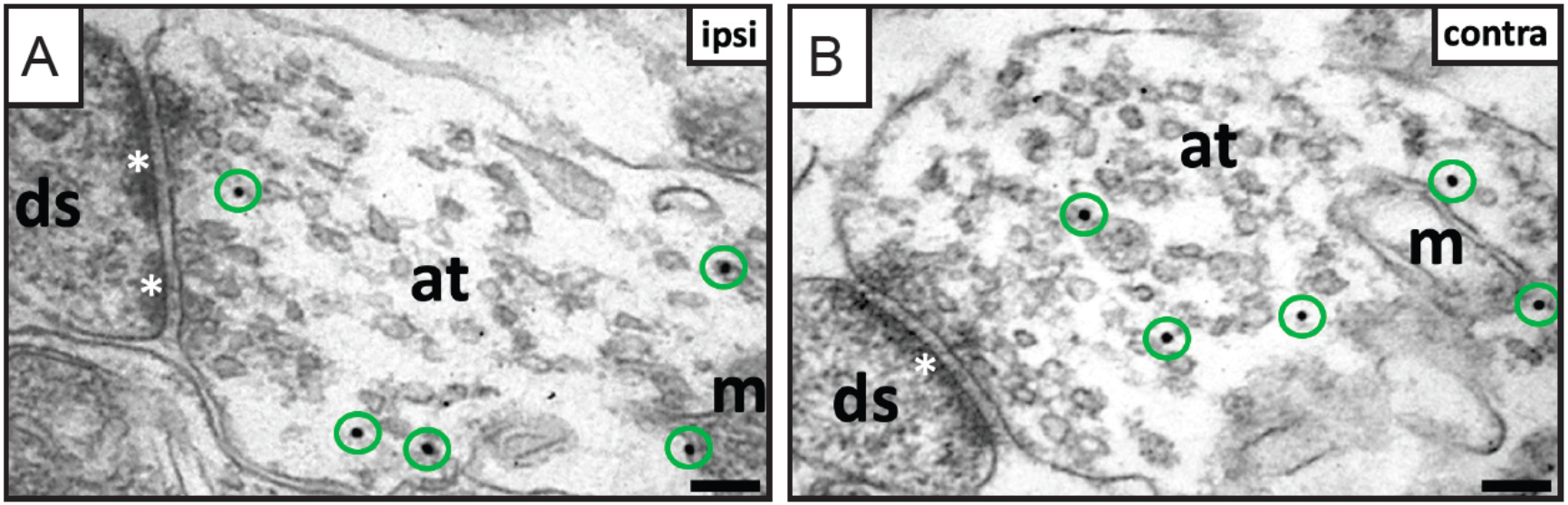
Grafted hES-iN establish functional synapses with host neurons in the stroke-injured rat brain. **A-B**, Asymmetric synapses with perforated (white asterisks, **A**) and continuous nonperforated postsynaptic densities (white asterisk, **B**) in host dendritic spines (ds) connected with grafted hES-iN derived-presynaptic axon terminals (at). Note: Gold particles are marked with green circles; m, mitochondrion. Scale bars, 0.1 μm.

## DISCUSSION

Here we demonstrate for the first time that neurons generated via direct programming of hES cells have the capacity to integrate into the stroke-injured rat brain already at 3 months after intracortical transplantation. Following only a 7-days NGN2-programming protocol *in vitro*, the hES cells gave rise to a homogeneous population of excitatory neurons characterized by co-expression of immature and mature neuronal markers and upper and deep cortical layer markers, and with electrophysiological properties of neurons in different stages of maturity. When grafting these cells into stroke-injured rat brain, hES-iNs survived and differentiated to mature, layer-specific cortical neurons. The hES-iNs sent widespread projections to different regions of the host brain, their axons became myelinated by host oligodendrocytes, and they established efferent synaptic connections with host neurons.

Several studies have demonstrated the potential of transplanted human cells, derived from ES and iPS cells, for replacing lost neurons following a cortical lesion in adult mouse or rat brain.^19,20^ The *in vitro* and *in vivo* generation of neurons via reprogramming of pluripotent stem cells has been demonstrated to be a rapid and efficient approach in terms of neuronal-related gene expression, morphology, connectivity, and electrophysiological properties.^21,22^ In most of previously published *in vitro* reprogramming protocols, addition of glial cells to the culture was needed to obtain mature functional neurons.^19,23^ However, in the absence of glia, Goparaju and collaborators found that transfection of human pluripotent stem cells with five mRNAs (encoding NGN1, NGN2, NGN3, ND1 and ND2) gives rise to neurons at day 7 of differentiation but only 50% of them showed 1 action potential^24^ In contrast, already after 7 days and without the presence of glial cells, our protocol generates neurons with capacity to fire multiple (65% of the total) or at least one action potential (35%), making our protocol faster than the ones described to date. However, it is important to take into account that full maturation of the neurons in culture will hamper their survival after transplantation. For this reason, an intermediate stage of maturation allowing both survival and fast maturation of the grafts should be ideal.

A problem when using pluripotent stem cells as a source for *in vivo* transplantation is the presence of cells with variable degree of maturation within the grafts.^25^ In our study, we found areas covered either by immature or mature neurons in the transplant. A small population of Sox2 positive cells was present in the cultures already prior to transplantation, and this presumed neural progenitor population could possibly be responsible for the occurrence of the immature areas in the grafts. In this context it should be mentioned that a transitory increase of Sox2 has been observed previously during *in vitro* programming of hES cells via overexpression of NGN2 between days 3 and 6.^19^ Some studies have shown that remaining undifferentiated neural precursor cells have a decreased proliferation capacity due to microenvironmental interactions with the host brain, decreasing the risk of teratoma formation^26,27^ Similar to our findings, previous studies have shown that human pluripotent stem cell-derived cortical progenitors transplanted into the injured adult visual cortex or somatosensory cortex, give rise to immature grafts, some of them displaying rosette-like structures and expressing proliferation markers 2 months after transplantation.^6,10^ These rosette-like structures were also observed in some areas of the graft in our transplantation setting. Also, in accordance to the presence of a minor population of neural progenitors among the transplanted cells, we found a small percentage of human astrocytes and oligodendrocytes within the graft.

We found that part of the DCX-expressing grafted cells co-expressed cortical markers, such as Satb2, which lead us to speculate that the immature grafted neurons present at 3 months after transplantation were already committed to become layer-specific neurons. In the mature areas of the transplant, we found a large number of grafted cells expressing markers from either upper (expressing Satb2 and Brn2) or deep (Ctip2 and Tbr1) cortical layers, organized into distinct clusters, suggesting some level of organization. This pattern of expression has been described previously after transplantation of fetal cortical cells into the damaged adult motor cortex.^28^

We have previously shown that cortically-primed iPS cell-derived neural progenitors, generated via small molecules and transplanted into a rat model of ischemic stroke, gave rise 77% DCX+ cells and 15% of NeuN+ cells 2 months after transplantation. About 10 % of the grafted cells were still proliferating at that time-point.^6^ At 6 months after transplantation, 40% of the cells were mature NeuN+ neurons while only 3% remained DCX positive. Similarly, when transplanting human pluripotent stem cell-derived cortical progenitors, derived using small molecules for cortical priming, into the injured adult visual cortex, a highly immature graft was observed at 2 months while the number of mature neurons had increased at 6 months.^10^ Our results showed that 35% of the graft area contained mature neurons already at 3 months post-transplantation. This finding provides suggestive evidence that programming by TF overexpression of hES cells gives rise to mature neural grafts in a shorter time after transplantation into the injured brain as compared to grafts generated by differentiation of human pluripotent stem cells using small molecules. These results on neuronal maturation are in line with the speed of axonal myelination. We observed that the hES-iN graft-derived axons were myelinated already at 3 months after transplantation whereas in previous studies with iPS-cell derived neurons generated via cortical priming with small molecules myelination was observed at 6 months. Importantly, at 3 months after transplantation into the somatosensory injured cortex, the hES-iNs gave rise to widespread projections the same areas (mainly ipsi-lesional cortex, corpus callosum and striatum) as observed with human iPS cell-derived neurons at 6 months post-grafting.^20^

The faster maturation of the intracortical grafts generated by TF-programming could possibly be explained by the higher expression of mature neuronal markers and the more mature electrophysiological properties of the neurons at the time of transplantation. Cortically primed iPS cell-derived neurons need 30 days of differentiation to exhibit molecular and electrophysiological properties similar to the ones observed in the present study in hES-iNs at day 7 of programming.^6,12^ It is important to highlight that different protocols for programming of neural cells can give rise to different cell populations. The overexpression of NGN2 in hES cells used here leads to the formation mainly of glutamatergic neurons, with a small proportion of astrocytes and oligodendrocytes within the graft. In contrast, generation of neural progenitors using a cortical priming protocol gives rise to glutamatergic neurons and oligodendrocytes in similar ratio and a small number of astrocytes.^8,9^

The present findings add another cell source to the repertoire of human-derived neural cells, generated from ES or iPS cells, with capacity to integrate morphologically and functionally in the stroke-injured brain after grafting. It should be emphasized, though, that the identification of several potentially useful cell sources for transplantation are only very early steps in a potential clinical translation. Much work remains, such as to generate the specific types of neurons and glia which will be crucial for reconstruction of specific stroke lesions in order to induce optimal functional recovery. Furthermore, even if efficient functional restoration can be obtained after xenografting in animal models, it is virtually unknown whether the transplanted neurons will reinstate appropriate connections and become properly myelinated also in the human *in vivo* setting. Recent findings, showing morphological and functional integration of grafted cortical neurons derived from pluripotent stem cells in organotypic cultures of adult human cortex^12^, provide supportive, promising evidence that this might be achievable in the future.

Taken together, hES-iN cells showed unique features in terms of the rapid generation of mature neurons both, *in vitro* and, after intracortical transplantation into a rat stroke model, giving rise to cortical layer-specific neurons able to integrate in the host injured circuitry within 3 months.

## AUTHOR CONTRIBUTIONS

S.P.-T., O.L., and Z.K. conceived the project. R.M.-C., M.H., L.J., N.A., O.T., E.M., B.C-S and S.P.-T. conducted the experiments and analyzed the data. R.M-C., S.P.-T., O.L., and Z.K. wrote the manuscript. R.M.-C., N.A., E.M., G.S., and O.T. were involved in collecting and/or assembly of data, data analysis, and interpretation. All authors reviewed and edited the manuscript.

## ACKNOWLEDGMENTS

We thank Daniel Capitán and Edita Lokmani for technical assistance. This work is supported by grants from the Swedish Research Council, Swedish Brain Foundation, Swedish Stroke Foundation, Regional Research Support from the Southern Swedish Healthcare Region, and the Swedish Government Initiative for Strategic Research Areas.

## REFERENCES

1. Poomalai G, Prabhakar S, Sirala Jagadesh N. Functional Ability and Health Problems of Stroke Survivors: An Explorative Study. Cureus. 2023.

2. De Celis-Ruiz E, Fuentes B, Moniche F, Montaner J, Borobia AM, Gutiérrez-Fernández M, Díez-Tejedor E. Allogeneic adipose tissue-derived mesenchymal stem cells in ischaemic stroke (AMASCIS-02): a phase IIb, multicentre, double-blind, placebo-controlled clinical trial protocol. BMJ Open. 2021;11(8):e051790.

3. Levy ML, Crawford JR, Dib N, Verkh L, Tankovich N, Cramer SC. Phase I/II Study of Safety and Preliminary Efficacy of Intravenous Allogeneic Mesenchymal Stem Cells in Chronic Stroke. Stroke. 2019;50(10):2835–2841.

4. Gong P, Zhang W, He Y, Wang J, Li S, Chen S, Ye Q, Li M. Classification and Characteristics of Mesenchymal Stem Cells and Its Potential Therapeutic Mechanisms and Applications against Ischemic Stroke. Yan J, ed. Stem Cells International. 2021;2021:1–13.

5. Law ZK, Tan HJ, Chin SP, Wong CY, Wan Yahya WNN, Muda AS, Zakaria R, Ariff MI, Ismail NA, Cheong SK, S Abdul Wahid SF, Mohamed Ibrahim N. The effects of intravenous infusion of autologous mesenchymal stromal cells in patients with subacute middle cerebral artery infarct: a phase 2 randomized controlled trial on safety, tolerability and efficacy. Cytotherapy. 2021;23(9):833–840.

6. Tornero D, Wattananit S, Grønning Madsen M, Koch P, Wood J, Tatarishvili J, Mine Y, Ge R, Monni E, Devaraju K, Hevner RF, Brüstle O, Lindvall O, Kokaia Z. Human induced pluripotent stem cell-derived cortical neurons integrate in stroke-injured cortex and improve functional recovery. Brain. 2013;136(12):3561–3577.

7. Tornero D, Tsupykov O, Granmo M, Rodriguez C, Grønning-Hansen M, Thelin J, Smozhanik E, Laterza C, Wattananit S, Ge R, Tatarishvili J, Grealish S, Brüstle O, Skibo G, Parmar M, et al. Synaptic inputs from stroke-injured brain to grafted human stem cell-derived neurons activated by sensory stimuli. Brain. 2017:aww347.

8. Palma-Tortosa S, Tornero D, Grønning Hansen M, Monni E, Hajy M, Kartsivadze S, Aktay S, Tsupykov O, Parmar M, Deisseroth K, Skibo G, Lindvall O, Kokaia Z. Activity in grafted human iPS cell–derived cortical neurons integrated in stroke-injured rat brain regulates motor behavior. Proceedings of the National Academy of Sciences. 2020;117(16):9094–9100.

9. Martinez-Curiel R, Jansson L, Tsupykov O, Avaliani N, Aretio-Medina C, Hidalgo I, Monni E, Bengzon J, Skibo G, Lindvall O, Kokaia Z, Palma-Tortosa S. Oligodendrocytes in human induced pluripotent stem cell-derived cortical grafts remyelinate adult rat and human cortical neurons. Stem Cell Reports. 2023:S2213671123001443.

10. Espuny-Camacho I, Michelsen KA, Linaro D, Bilheu A, Acosta-Verdugo S, Herpoel A, Giugliano M, Gaillard A, Vanderhaeghen P. Human Pluripotent Stem-Cell-Derived Cortical Neurons Integrate Functionally into the Lesioned Adult Murine Visual Cortex in an Area-Specific Way. Cell Reports. 2018;23(9):2732–2743.

11. Jgamadze D, Lim JT, Zhang Z, Harary PM, Germi J, Mensah-Brown K, Adam CD, Mirzakhalili E, Singh S, Gu JB, Blue R, Dedhia M, Fu M, Jacob F, Qian X, et al. Structural and functional integration of human forebrain organoids with the injured adult rat visual system. Cell Stem Cell. 2023;30(2):137–152.e7.

12. Grønning Hansen M, Laterza C, Palma-Tortosa S, Kvist G, Monni E, Tsupykov O, Tornero D, Uoshima N, Soriano J, Bengzon J, Martino G, Skibo G, Lindvall O, Kokaia Z. Grafted human pluripotent stem cell-derived cortical neurons integrate into adult human cortical neural circuitry. STEM CELLS Translational Medicine. 2020:sctm.20-0134.

13. Lindvall O. Treatment of Parkinson’s disease using cell transplantation. Philosophical Transactions of the Royal Society B: Biological Sciences. 2015;370(1680):20140370.

14. Miskinyte G, Grønning Hansen M, Monni E, Lam M, Bengzon J, Lindvall O, Ahlenius H, Kokaia Z. Transcription factor programming of human ES cells generates functional neurons expressing both upper and deep layer cortical markers. Zheng JC, ed. PLOS ONE. 2018;13(10):e0204688.

15. Chen ST, Hsu CY, Hogan EL, Maricq H, Balentine JD. A model of focal ischemic stroke in the rat: reproducible extensive cortical infarction. Stroke. 1986;17(4):738–743.

16. Oki K, Tatarishvili J, Wood J, Koch P, Wattananit S, Mine Y, Monni E, Tornero D, Ahlenius H, Ladewig J, Brüstle O, Lindvall O, Kokaia Z. Human-Induced Pluripotent Stem Cells form Functional Neurons and Improve Recovery After Grafting in Stroke-Damaged Brain. STEM CELLS. 2012;30(6):1120–1133.

17. Renier N, Wu Z, Simon DJ, Yang J, Ariel P, Tessier-Lavigne M. iDISCO: A Simple, Rapid Method to Immunolabel Large Tissue Samples for Volume Imaging. Cell. 2014;159(4):896–910.

18. Schneggenburger R, Sakaba T, Neher E. Vesicle pools and short-term synaptic depression: lessons from a large synapse. Trends in Neurosciences. 2002;25(4):206– 212.

19. Zhang Y, Pak C, Han Y, Ahlenius H, Zhang Z, Chanda S, Marro S, Patzke C, Acuna C, Covy J, Xu W, Yang N, Danko T, Chen L, Wernig M, et al. Rapid Single-Step Induction of Functional Neurons from Human Pluripotent Stem Cells. Neuron. 2013;78(5):785– 798.

20. Buchlak QD, Esmaili N, Moore J. Opportunities for developing neural stem cell treatments for acute ischemic stroke: A systematic review and gap analysis. Journal of Clinical Neuroscience. 2024;120:64–75.

21. Clark IH, Roman A, Fellows E, Radha S, Var SR, Roushdy Z, Borer SM, Johnson S, Chen O, Borgida JS, Steevens A, Shetty A, Strell P, Low WC, Grande AW. Cell Reprogramming for Regeneration and Repair of the Nervous System. Biomedicines. 2022;10(10):2598.

22. Bocchi R, Masserdotti G, Götz M. Direct neuronal reprogramming: Fast forward from new concepts toward therapeutic approaches. Neuron. 2022;110(3):366–393.

23. Nehme R, Zuccaro E, Ghosh SD, Li C, Sherwood JL, Pietilainen O, Barrett LE, Limone F, Worringer KA, Kommineni S, Zang Y, Cacchiarelli D, Meissner A, Adolfsson R, Haggarty S, et al. Combining NGN2 Programming with Developmental Patterning Generates Human Excitatory Neurons with NMDAR-Mediated Synaptic Transmission. Cell Reports. 2018;23(8):2509–2523.

24. Goparaju SK, Kohda K, Ibata K, Soma A, Nakatake Y, Akiyama T, Wakabayashi S, Matsushita M, Sakota M, Kimura H, Yuzaki M, Ko SBH, Ko MSH. Rapid differentiation of human pluripotent stem cells into functional neurons by mRNAs encoding transcription factors. Scientific Reports. 2017;7(1):42367.

25. Ford E, Pearlman J, Ruan T, Manion J, Waller M, Neely GG, Caron L. Human Pluripotent Stem Cells-Based Therapies for Neurodegenerative Diseases: Current Status and Challenges. Cells. 2020;9(11):2517.

26. Bernal A, Arranz L. Nestin-expressing progenitor cells: function, identity and therapeutic implications. Cellular and Molecular Life Sciences. 2018;75(12):2177–2195.

27. Vonderwalde I, Azimi A, Rolvink G, Ahlfors J-E, Shoichet MS, Morshead CM. Transplantation of Directly Reprogrammed Human Neural Precursor Cells Following Stroke Promotes Synaptogenesis and Functional Recovery. Translational Stroke Research. 2020;11(1):93–107.

28. Ballout N, Frappé I, Péron S, Jaber M, Zibara K, Gaillard A. Development and Maturation of Embryonic Cortical Neurons Grafted into the Damaged Adult Motor Cortex. Frontiers in Neural Circuits. 2016;10.

