## Supplementary Figures and Tables for "Human cortical neurons rapidly generated by direct ES cell programming integrate into stroke-injured rat cortex"

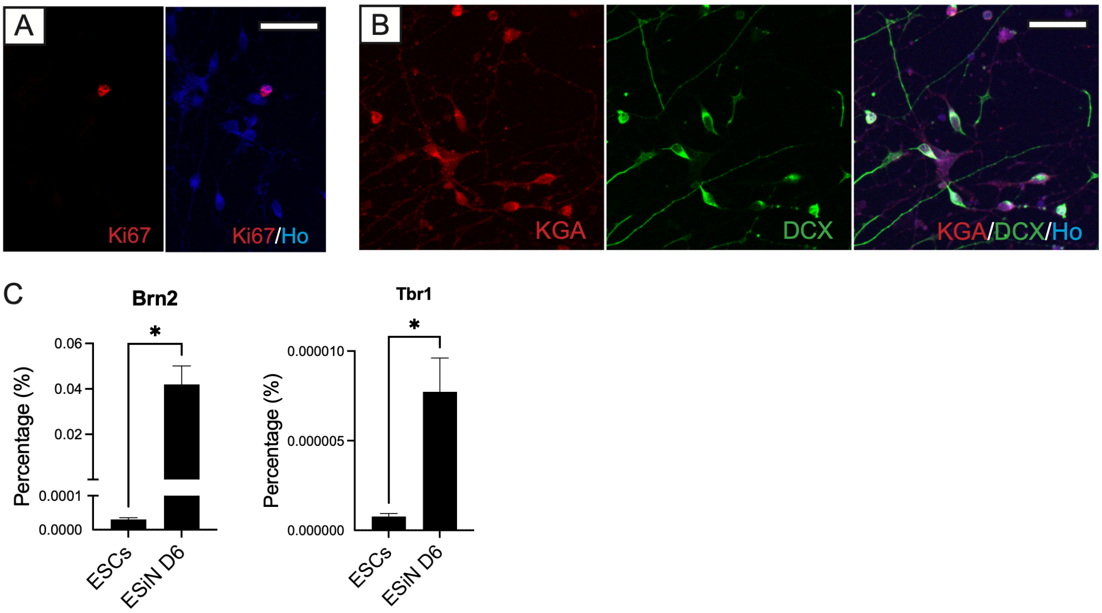


**Figure S1**. ***In vitro* programming of hES cells give rise to glutamatergic cortical neurons**. **A, B**, Representative confocal images showing the proliferation marker Ki67 (**A**), the glutamatergic marker KGA and the immature neuronal marker DCX (**B**). Scale bar, 50 μm. **C**, Gene expression of *BRN2* and *TBR1* in hES-iNs after 6 days of programming. Data are presented as mean ± SEM. Significance was set as $p < 0.05$.

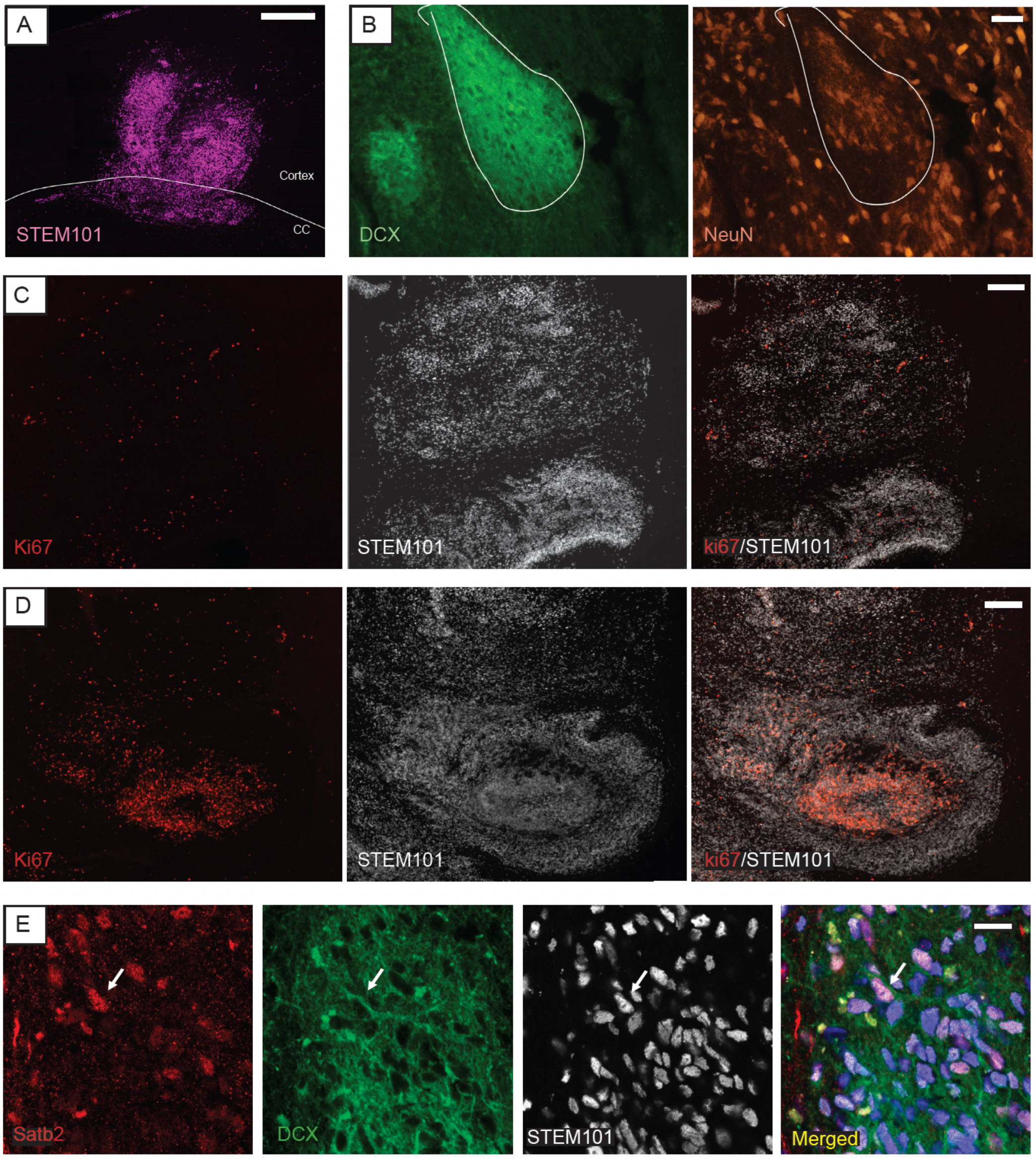


**Figure S2. Intracortical transplantation of hES-iNs give rise to heterogeneous grafts in terms location and proliferation**. **A**, Overview of a large graft, identified by STEM101, showing grafted hES-iNs in the cortex and corpus callosum 3 months after transplantation in the injured-stroke cortex. Scale bar, 500 µm. **B**, hES-iNs show areas of colocalization of the immature neuronal marker DCX and mature neuronal marker NeuN (white outline). Scale bar, 20 µm. **C-D**, Proliferative grafted hES-iNs shown by human nuclear marker STEM101 and proliferation marker ki67. **C**, Graft with sparse grafted derived proliferative cells. **D**, Grafted hES-iNs higher proliferation in less mature areas identified by rosette morphology. Scale bar, 200 µm. **E**, Representative confocal image showing the immature neuronal marker DCX and the upper cortical layer marker Satb2 colocalizing with STEM101. Arrow indicate colocalization. Nuclear staining (Ho: Hoechst, blue) is included in merged panel. Scale bar, 20 µm.

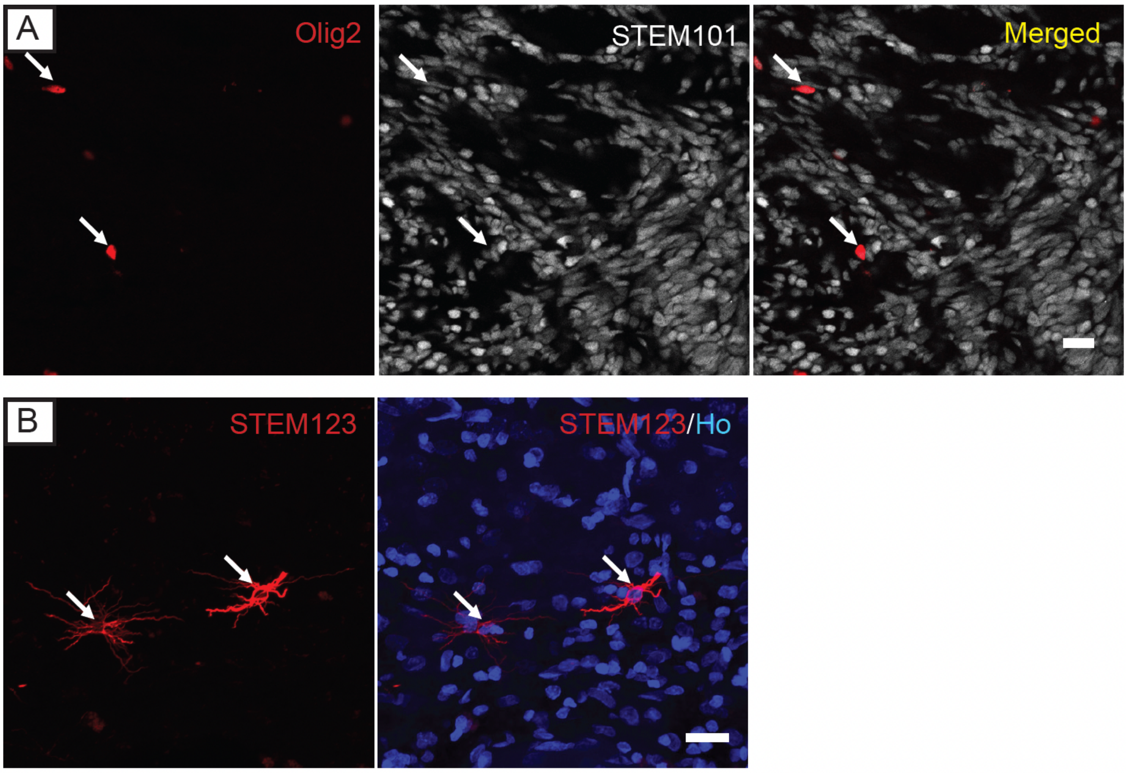


**Figure S3**. **Grafted hES-iNs give rise to a small population of oligodendrocytes and astrocytes 3 months after transplantation in the injured-stroke cortex**. **A-B**, Representative confocal images showing graft-derived oligodendrocytes shown by linage oligodendrocyte marker Olig2 (**A**) and graft-derived astrocytes shown by specific human GFAP marker, STEM123 (**B**). Arrows indicate colocalization. Scale bar, 20 μm.

| Antibody | Host species | *In vitro* | *In vivo* | Notes | Company |
|---|---|---|---|---|---|
| Primary antibodies | | | | | |
| Brn2 | Goat | 1:400 | 1:400 | AR | Santa Cruz |
| Ctip2 | Rabbit | 1:100 | 1:200 | AR | Sigma |
| DCX | Goat | 1:400 | 1:400 | | Santa Cruz |
| KGA | Rabbit | 1:1000 | - | | Abcam |
| Ki67 | Rabbit | 1:500 | 1:250 | | Abcam |
| Nanog | Rabbit | 1:150 | - | | Abcam |
| NeuN | Rabbit | 1:500 | 1:500 | | Abcam |
| Olig2 | Rabbit | 1:500 | 1:500 | | Abcam |
| Satb2 | Mouse | 1:100 | - | AR | Abcam |
| Satb2 | Rabbit | - | 1:200 | AR | Abcam |
| STEM101 | Mouse | - | 1:500 | | StemCells |
| STEM121 | Mouse | - | 1:500 | | StemCells |
| STEM123 | Mouse | - | 1:2000 | | StemCells |
| Sox2 | Rabbit | 1:200 | 1:200 | | Merk Millipore |
| Tbr1 | Rabbit | - | 1:300 | | Gifted |
| Tuj1 | Mouse | 1:2000 | - | | Sigma |
| Secondary Antibodies | | | | | |
| 488 anti-Goat | Donkey | 1:500 | 1:500 | | Jackson ImmunoResearch |
| 488 anti-Mouse | Donkey | 1:500 | 1:500 | | Jackson ImmunoResearch |
| 488 anti-Rabbit | Donkey | 1:500 | 1:500 | | Jackson ImmunoResearch |
| Cy3 anti-Goat | Donkey | 1:500 | - | | Jackson ImmunoResearch |
| Cy3 anti-Mouse | Donkey | 1:500 | 1:500 | | Jackson ImmunoResearch |
| Cy3 anti-Rabbit | Donkey | 1:500 | 1:500 | | Jackson ImmunoResearch |
| 647 anti-Mouse | Donkey | 1:500 | 1:500 | | Jackson ImmunoResearch |
| 647 anti-Rabbit | Donkey | 1:500 | 1:500 | | Jackson ImmunoResearch |

**Table S1.** List of primary and secondary antibodies. **Related to section: *Immunostainings and quantifications***.

**VIDEO SUPPLEMENTARY 4. Grafted hES-iN send widespread projections to different brain regions in 3 months after transplantation into the injured rat brain.** Grafted cells immunostained with human nuclear marker STEM101 and human cytoplasmic marker STEM121.
